# A Compensatory RNase E Variation Increases Iron Piracy and Virulence in Multidrug-Resistant *Pseudomonas aeruginosa* during Macrophage Infection

**DOI:** 10.1101/2022.10.24.513460

**Authors:** Mylene Vaillancourt, Anna Clara Milesi Galdino, Sam P. Limsuwannarot, Diana Celedonio, Elizabeth Dimitrova, Matthew Broerman, Catherine Bresee, Yohei Doi, Janet S. Lee, William C. Parks, Peter Jorth

## Abstract

During chronic cystic fibrosis (CF) infections, evolved *Pseudomonas aeruginosa* antibiotic resistance is linked to increased pulmonary exacerbations, decreased lung function, and hospitalizations. However, the virulence mechanisms underlying worse outcomes caused by antibiotic resistant infections are poorly understood. Here, we investigated evolved aztreonam resistant *P. aeruginosa* virulence mechanisms. Using a macrophage infection model combined with genomic and transcriptomic analyses, we show that a compensatory mutation in the *rne* gene, encoding RNase E, increased siderophore gene expression, causing macrophage ferroptosis and lysis. Macrophage killing could be eliminated by treatment with the iron mimetic gallium. RNase E variants were abundant in clinical isolates, and CF sputum gene expression data show that clinical isolates phenocopied RNase E variant functions during macrophage infection. Together these data show how *P. aeruginosa* RNase E variants can cause host damage via increased siderophore production and host cell ferroptosis but may also be targets for gallium precision therapy.

## Introduction

*Pseudomonas aeruginosa* is a ubiquitous bacterium that thrives in a wide range of environments including water, soils, plants, and animal tissues ^1^. In clinical settings, *P. aeruginosa* is one of the prevailing pathogens involved in acute nosocomial infections and chronic lung infections in cystic fibrosis (CF) ^2^. During acute infection by *P. aeruginosa*, innate immune cells are critical for clearance by rapidly decreasing bacterial loads during the initial infection, even before the first administration of antibiotics 3. In particular, macrophages are the predominant immune cells to phagocytose *P. aeruginosa* after bacterial inhalation ^4^. More importantly, bacterial clearance by macrophages prevents neutrophil infiltration and subsequent inflammation ^4–6^. During lung infection, recruitment of peripheral monocytes/macrophages is essential for bacterial clearance and reduced neutrophilia ^5^. In the same line, lower numbers of activated monocytes/macrophages are associated with the presence of *P. aeruginosa* infection and disease severity in CF ^7^. These observations underscore a crucial role for macrophages in bacterial clearance and homeostasis of the lung.

Bacteria evolve several mechanisms to escape phagocytosis and colonize the lung environment. For instance, *P. aeruginosa* use rhamnolipids, a glycolipid biosurfactant, to break through epithelial barriers and escape phagocytosis by immune cells ^8^. Virulent strains also use the type III secretion system (T3SS) to induce macrophage lysis ^9,10^. Bacteria can use molecules from lysed host cells for their own benefits. For example, *P. aeruginosa* secretes siderophores that scavenge iron from host cells and are recaptured by the bacteria for their metabolic activities ^11^.

*P. aeruginosa* is also able to maintain its growth using anaerobic respiration by denitrification in the CF lung and sputum ^12^. While these phenotypes undoubtedly benefit *P. aeruginosa* during infection, the importance of these different mechanisms can be unclear due to genotypic and phenotypic diversification ^13,14^. Such diversification can lead to the overexpression or genotypic loss of numerous virulence systems, metabolic versatility, and antibiotic resistance ^15–17^. As a result, despite extensive work in this field, the response of *P. aeruginosa* to different environmental triggers and the downstream mechanisms involved in virulence and lung adaptation remain elusive.

During chronic lung infection, pathogens experience selective pressure from intense antibiotic therapies ^18,19^. Over the past decades, extensive use of antibiotics has led to increased prevalence of multidrug- and extensively drug-resistant *P. aeruginosa* worldwide ^20–22^. Although the acquisition of antibiotic resistance may be correlated with decreased virulence and fitness, multiple studies found associations between multidrug-resistant *P. aeruginosa* infections and pulmonary exacerbations, decreased lung function, and hospitalizations in people with CF ^2,23^. In line with these clinical observations, multidrug-resistant *P. aeruginosa* strains can exhibit virulence evolved from either the resistance mutation or various types of second-site compensatory mutations ^24–27^. However, mechanisms by which antibiotic resistance leads to hypervirulence are poorly understand.

Our group recently showed that a loss-of-function mutation in the *mexR* gene, which led to aztreonam resistance by overexpressing MexAB-OprM efflux pump, was sufficient to increase bacterial virulence *in vivo* ^27^. This strain also displayed increased swarming motility resulting from a second-site mutation impacting the MexEF-OprN efflux pump ^27^. This second-site mutation enhanced virulence *in vivo*, independently of *mexR* and MexAB-OprM. These data highlight the unexpected dangers of antibiotic selection for hypervirulent bacteria and the urgent need to find alternative therapies to eradicate these “superbugs”.

In the present study, we sought to investigate the virulence mechanisms of the *P. aeruginosa* AzEvC10 mutant, a PAO1 strain that was evolved under cyclic aztreonam exposure ^26^. This strain first evolved a mutation in the *nalD* gene, resulting in overexpression of the MexAB-OprM efflux pump and multidrug resistance. This mutant was also hypervirulent *in vivo* compared to wild-type (WT) PAO1. However, it was unclear whether the AzEvC10 strain’s virulence was dependent upon the *nalD* mutation or a second-site compensatory mutation. Furthermore, the virulence mechanisms upregulated in the AzEvC10 strain remained elusive. Here we determine mechanisms of hypervirulence of the AzEVC10 evolved strain and ask whether this virulence is caused directly by its antibiotic resistance mutation, or via a second-site compensatory mutation. We also determine the host cell types directly affected by this strain’s increased virulence, identify a potential treatment to kill these hypervirulent *P. aeruginosa* variants, and reveal similarities between this lab-evolved strain and *P. aeruginosa* clinical isolates.

## Results

### Aztreonam-evolved *P. aeruginosa* replicates more during macrophage infection

To characterize the virulence of *P. aeruginosa* AzEvC10 mutant, we assessed if there was a difference in clearance between the mutant and WT PAO1 by infecting bone marrow-derived macrophages (BMDM), BM-derived neutrophils, or airway epithelial cells (AEC) with 2.5 × 10^7^ CFU/ml of either strain. At 6 h post infection (hpi) of BMDM, the AzEvC10 mutant had increased in number by about 1 log_10_ whereas the WT parental strain showed no change from the initial inoculum (*p*< 0.01) (Figure 1A). In addition, the AzEvC10 mutant induced higher cytotoxicity of BMDM (44% ± 8.2) compared to the WT strain (20% ± 5.2) (*p*<0.05) (Figure 1B). These differences were specific to BMDM, as there was no difference in the replication of (Figure 1A) or cytotoxicity (data not shown) by the two strains in BM-derived neutrophils and AEC.

**Figure 1.**
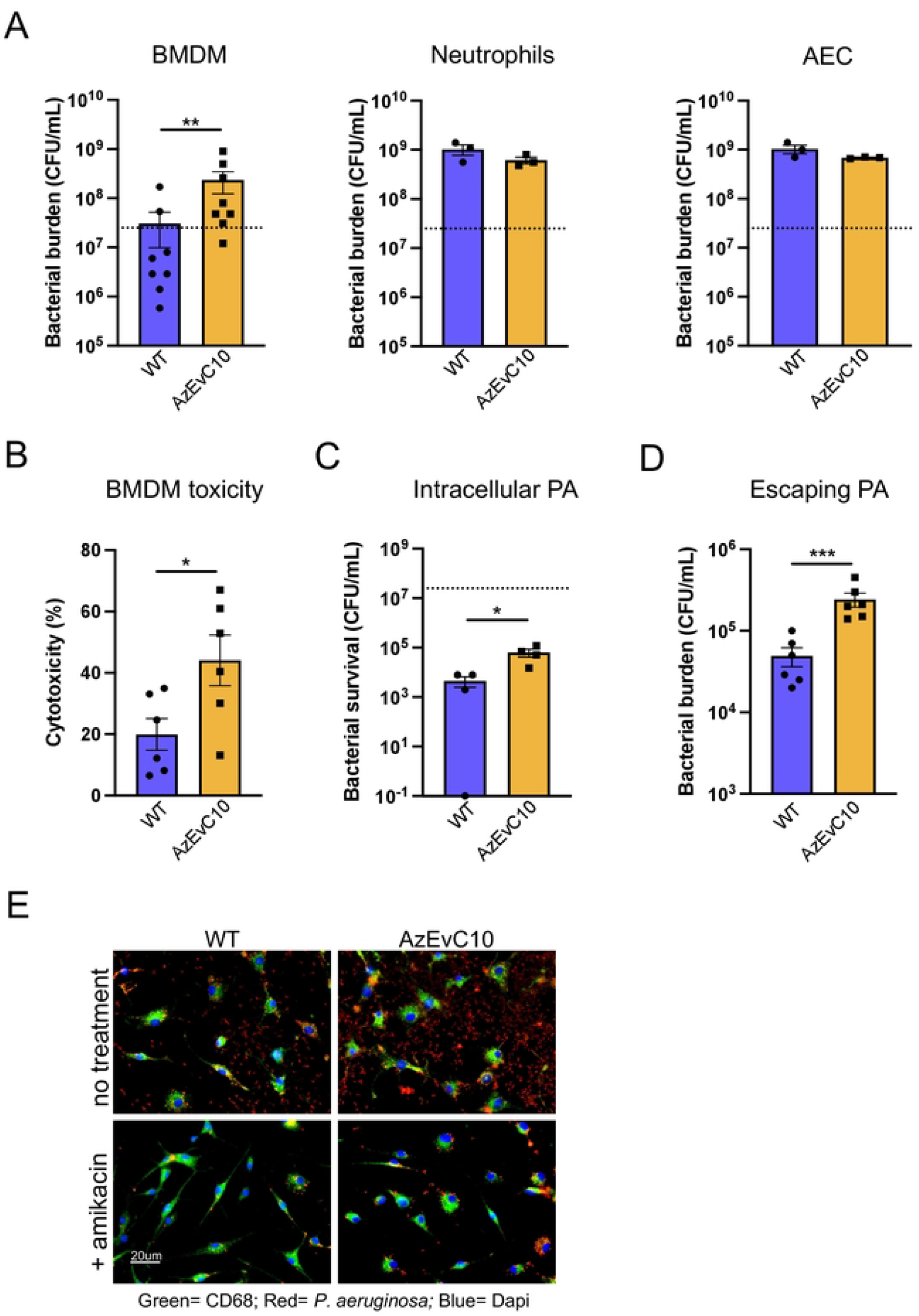
An aztreonam-evolved *P. aeruginosa* mutant replicates more during macrophage infection. **A**. BMDM, BM-derived neutrophil, or AEC cells were infected with WT PAO1 or the AzEvC10 mutant for 6 h. Dotted lines represent the initial infection inoculum, 2.5 × 10^7^ CFU/mL. Bacterial burden at 6 hpi was determined by viable CFU plate counts. Analysis was matched for experimental repeat. **B**. BMDM cytotoxicity assessed by LDH assay at 6 hpi by WT or AzEVC10. Analysis was matched for experimental repeat. **C**. Intracellular survival of bacteria in BMDM at 6 hpi, following amikacin treatment at 1 hpi for a total of 5 h to kill extracellular bacteria. Viable intracellular bacteria were determined by CFU plate counts. **D**. Escaping bacteria at 6 hpi: Amikacin was added in media at 1 hpi and left for 1 h. Then cells were incubated in fresh media (no amikacin) for another 4 h. Escaping bacteria were determined by CFU plate counts. **E**. Images of BMDM infections at 6 hpi by immunofluorescence showing macrophages in green, *P. aeruginosa* in red, and DAPI in blue. n = 3-6 independent replicates for each experiment. \**p*<0.05, #x002A;\**p*<0.01, \*\*\**p*<0.001, \*\*\*\**p*<0.0001. See Table S5 for statistical tests used and exact *p*-values.

We tested if increased proliferation of the AzEvC10 mutant was due its capacity to use growth medium components as a source of carbon and energy. However, we found that both strains had net lethality in host cell-free medium but that there was a significant survival difference between the strains. At 6 hpi in cell-free medium, the number of viable WT PAO1 was 3.5 log_10_ less than the initial inoculum, whereas the number of viable AzEvC10 mutants was reduced only 0.8 log_10_ (Figure S1A). These findings demonstrate that the AzEvC10 mutant has acquired a significant survival advantage – nearly 1,000-fold better than WT – in a nutrient restrictive environment.

We also assessed if the survival and subsequent escape of phagocytosed bacteria contributed to the increased bacterial burden of AzEvC10 mutant in macrophage cultures. To test this, we infected BMDM with the mutant and WT and 1 h later added 200 μg/mL amikacin, an antibiotic that does not enter host cells, to kill extracellular bacteria. This concentration of amikacin effectively killed both WT PAO1 and AzEvC10 mutant bacteria (Figure S1B). At 6 hpi (i.e., 5 h post-amikacin), the number of surviving intracellular AzEvC10 mutants was significantly greater (about 1 log_10_) than the number of intracellular viable WT PAO1 (*p*<0.05) (Figure 1C) and this could be due to either higher phagocytosis of mutant bacteria by macrophages, increased mutant bacterial proliferation inside host cells, or increased intracellular survival. Regardless, to determine if these intracellular surviving bacteria could escape from BMDM and proliferate extracellularly, we treated BMDM medium with amikacin at 1 hpi, washed the cells, and added fresh medium without antibiotic. Compared to WT infected cells, we found that extracellular bacterial counts were about 1 log_10_ increased in AzEvC10 mutant infected cultures at 6 hpi (Figure 1D, 1E), proportional to the difference in viable intracellular bacteria (Figure 1C). These data suggest that both WT PAO1 and AzEvC10 mutant can escape BMDMs and proliferate extracellularly with similar efficiency. Although the specific cause of the increased intracellular bacterial recovery needs to be defined, overall, our studies indicate that AzEvC10 adapts better than WT PAO1 to the host environment and that this growth advantage was enhanced by the presence of macrophages. Below we explore the bacterial and host pathways involved in the mutant’s growth advantage.

### AzEvC10 mutant hypervirulence is caused by a compensatory mutation in RNase E

The aztreonam-evolved AzEvC10 mutant first acquired a loss-of-function point mutation in the *nalD* gene (Figure S2A) in response to the aztreonam selective pressure, which caused overexpression of the MexAB-OprM efflux pump and a multidrug resistance phenotype ^26^. Recently, we found that mutation of another *mexAB-oprM* regulator was sufficient to increase *P. aeruginosa* virulence ^27^. Thus, we hypothesized that the AzEVC10 mutant’s virulence was being driven by its *nalD* mutation. To test this, we complemented a functional wild-type *nalD* gene in the AzEvC10 mutant and infected BMDM to assess if this would reverse the hypervirulence. Complementation of the *nalD* gene was confirmed by mRNA expression quantification and aztreonam MIC (Figure S2B, S2C). Unexpectedly, *nalD* complementation in AzEvC10 did not reverse bacterial growth and cytotoxicity during BMDM infection (Figure S2D, S2E). This was further confirmed when we infected BMDM with mutants in which *mexAB* genes were deleted (Figure S2F-S2I).

To explore if the AzEvC10 mutant’s hypervirulence could be explained by a second-site compensatory mutation, we used whole genome sequencing and found that the strain evolved a 50-bp deletion (bases 3121-3170/3174) in the *rne* gene (Figure 2A, 2B) encoding RNase E, an endonuclease involved in RNA turnover. Because compensatory mutations can restore fitness and/or virulence, and due to the nature of RNase E activity, we hypothesized that the *rne* variation was involved in the virulence and growth of the AzEvC10 mutant during macrophage infection. To begin testing the virulence consequences of the *rne* mutation, we created single *nalD*_T158P_ and *rne*_Δ50bp_ mutants in a wild-type *P. aeruginosa* background and used an *rne* mutant containing a transposon inserted at the position 2945 of the gene (*rne*::Tn) ^28^. We also reversed the *rne* mutation directly in the AzEvC10 chromosome at its native position (AzEvC10 *rne*_WT_) as a rescue. As expected, the aztreonam resistance seen in AzEvC10 mutant was caused by the mutation in *nalD* gene (Figure 2C). However, we found that bacterial burden during BMDM infection and BMDM cytotoxicity were increased in AzEvC10, *rne*_Δ50bp_, and *rne*::Tn mutants compared to WT, *nalD*_T158P_, and AzEvC10 *rne*_WT_ strains (Figure 2D-2F). These results confirmed the role of the RNase E variant in the hypervirulent phenotype and growth of AzEvC10 mutant during macrophage infection.

**Figure 2.**
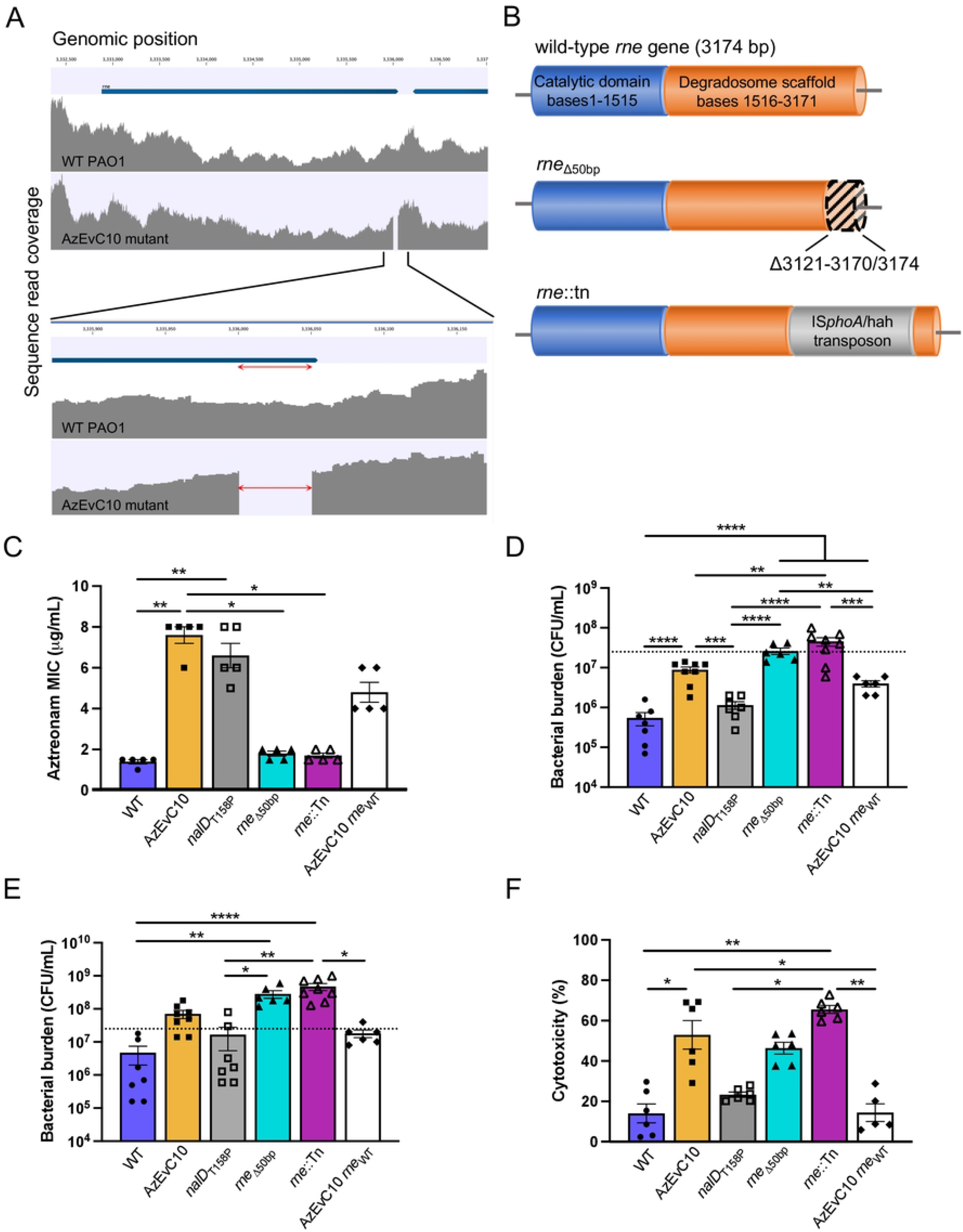
AzEvC10 mutant hypervirulence is caused by an RNase E mutation. **A**. Genome diagram showing coverage of DNA sequencing reads aligning to the *rne* gene. The deletion of bases 3121-3170/3174 nt of the *rne* gene in the AzEvC10 mutant are enlarged in the bottom panel. Genome coverage plots generated from sequencing read alignments to the PAO1 reference genome are indicated in grey. **B**. Schematic representation of the *rne* gene and corresponding RNase E protein domain description. WT PAO1, *nalD*_T158P_, and AzEvC10 *rne*_WT_ mutants carry a wild-type *rne* gene. AzEvC10 and *rne*_Δ50bp_ mutants have a deletion of the bases 3121-3170/3174 nt in the *rne* gene. The *rne*::Tn mutant carries an IS*phoA*/hah transposon inserted at the position 2945 nt of the gene. **C**. Aztreonam MIC was assessed by Etest for the indicated bacteria. **D-E**. BMDM cells were infected with WT PAO1 or different *nalD* and *rne* mutants at MOI:100 for 3 h (**D**) and 6 h (**E**). Bacterial burden was determined by viable CFU plate counts. Dotted lines represent the initial infection inoculum, 2.5 × 10^7^ CFU/mL. **F**. BMDM cytotoxicity was assessed by LDH assay at 6 hpi with indicated bacteria. n=5-8 independent replicates for each experiment \**p*<0.05, \*\**p*<0.01, \*\*\**p*<0.001, \*\*\*\**p*<0.0001. See Table S5 for statistical tests used and exact *p*-values.

### Iron acquisition genes are upregulated in the AzEvC10 mutant during BMDM infection

While the *rne* variation was responsible for the increased virulence of the AzEvC10 mutant, the mechanisms driving the *rne* variant’s increased virulence were unclear. To understand the mechanisms underlying the AzEvC10 mutant phenotype, we infected BMDM and performed RNA sequencing at 3 hpi and 6 hpi. We first compared the AzEvC10 mutant gene expression to WT PAO1 during the BMDM infections at the two timepoints. We found a marked upregulation of the two *P. aeruginosa* siderophore biogenesis pathways at both 3 hpi and 6 hpi in the AzEvC10 mutant (Figure 3A, 3B). These pathways include genes involved in pyoverdine biosynthesis (*pvdA, pvdD, pvdE, pvdF, pvdG, pvdH, pvdJ, pvdL, pvdN,pvdO, pvdP, pvdQ*), transport (*pvdR, pvdT, opmQ*), and recycling (*fpvA*). Genes encoding the pyochelin biosynthesis enzymes (*pchABCD, pchEFG*), receptors (*fptA, fptB*), and their transcriptional regulator (*pchR*) were also upregulated at 3 hpi and even more so at 6 hpi. Upregulation of heme acquisition genes (*hasAp, hasD, hasE and hasR*) was observed but only at 6 hpi. Interestingly, none of these pathways were upregulated when we grew the strains in LB broth (Figure 3B), suggesting that this mutant’s response was specific to the macrophage infection.

**Figure 3.**
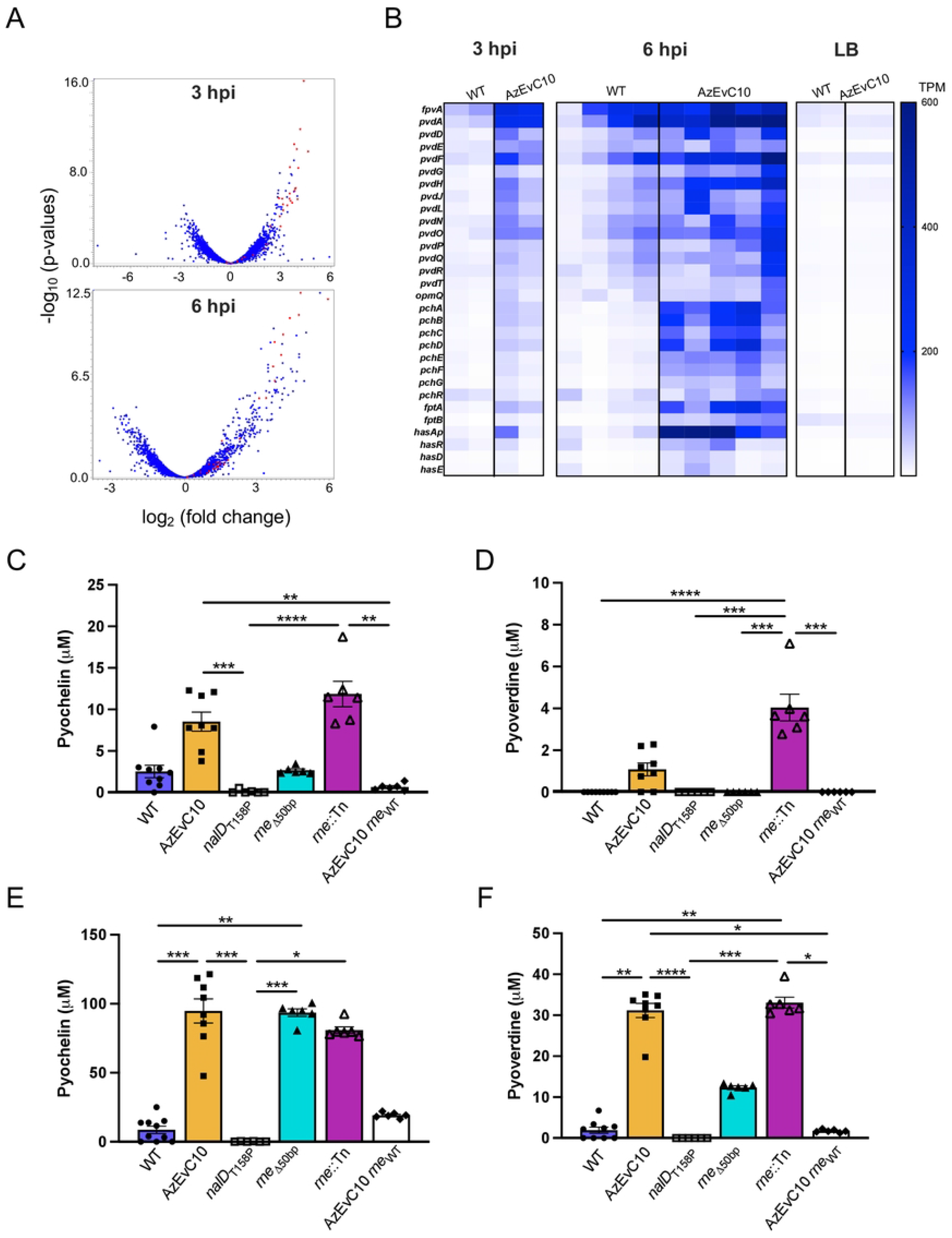
Iron acquisition genes are upregulated in the AzEvC10 mutant during macrophage infection. **A-B**. Genes involved in iron acquisition genes were upregulated in the AzEvC10 mutant compared to WT PAO1 during BMDM infection at 3 hpi (n=2) and 6 hpi (n=4-5), but not when grown in LB broth (n=2). **A**. Volcano plot highlighting differentially expressed iron acquisition genes in red during BMDM infection. **B**. Heat map indicates expression of indicated genes in individual biological replicates expressed in normalized number of reads (TPM) during WT and AzEvC10 BMDM infections at indicated time points. **C-F**. Pyochelin and pyoverdine production by given strains measured by fluorescence at 3 hpi **(C-D)** and 6 hpi **(E-F)** and quantified using standard curves. n=5-8 independent replicates for each experiment. \**p*<0.05, \*\**p*<0.01, \*\*\**p*<0.001, \*\*\*\**p*<0.0001. See Table S5 for statistical tests used and exact *p*-values.

To confirm the effect of the RNase E variation on increased siderophore biogenesis, we measured pyochelin and pyoverdine secreted in the growth medium during BMDM infection. Both siderophores were barely detectable in WT, *nalD*_T158P_, and AzEvC10 *rne*_WT_ strains at 3 hpi and 6 hpi (Figure 3C-3F). In contrast, the AzEvC10 and *rne*::Tn mutants had produced detectable pyochelin and pyoverdine by 3 hpi, and at 6 hpi, the concentrations increased further to 94 ± 24 μM and 80 ± 5 μM, respectively, for pyochelin (Figure 3E) and 31 ± 5 μM and 33 ± 3 μM, respectively, for pyoverdine (Figure 3F). Surprisingly, the *rne*_Δ50bp_ mutant did not secrete siderophores at 3 hpi, as seen in AzEvC10 and *rne*::Tn mutants. However, the mutant did secrete pyochelin (93 ± 7 μM) and pyoverdine (12 ±1 μM) at 6 hpi. While the reason for this difference is not clear, it is possible that the dynamics of increased siderophore production vary slightly based on genetic background and the nature of the *rne* variation.

### The AzEvC10 mutant carries specific gene expression signatures during BMDM infection

It was unclear if the increased iron acquisition capacity in the AzEvC10 mutant preceded or followed its increased proliferation during BMDM infection. This distinction is important to resolve because either: 1) the increased capacity to capture iron could improve bacterial fitness under stressful conditions such as being in contact with host immune cells, or 2) increased mutant proliferation could create a greater need for iron and trigger the biosynthesis of pyochelin and pyoverdine siderophores. To resolve these two possibilities, we first performed Gene Ontology Enrichment Analyses to identify processes that were differentially regulated by the AzEvC10 mutant at 3 hpi and 6 hpi relative to WT infections. Our analyses revealed two distinct gene expression patterns between the two timepoints. At 3 hpi, the AzEvC10 mutant showed increased gene expression of siderophore biosynthesis and secretion processes, including siderophore and pyoverdine metabolic processes, T3SS, cell projection and assembly, protein membrane transport and secretion (Figure 4A). In contrast, at 6 hpi, genes in metabolic pathways such as isoprenoid, lipid, terpene carboxylic acid, and amino acid catabolism were differentially expressed (Figure 4B).

**Figure 4.**
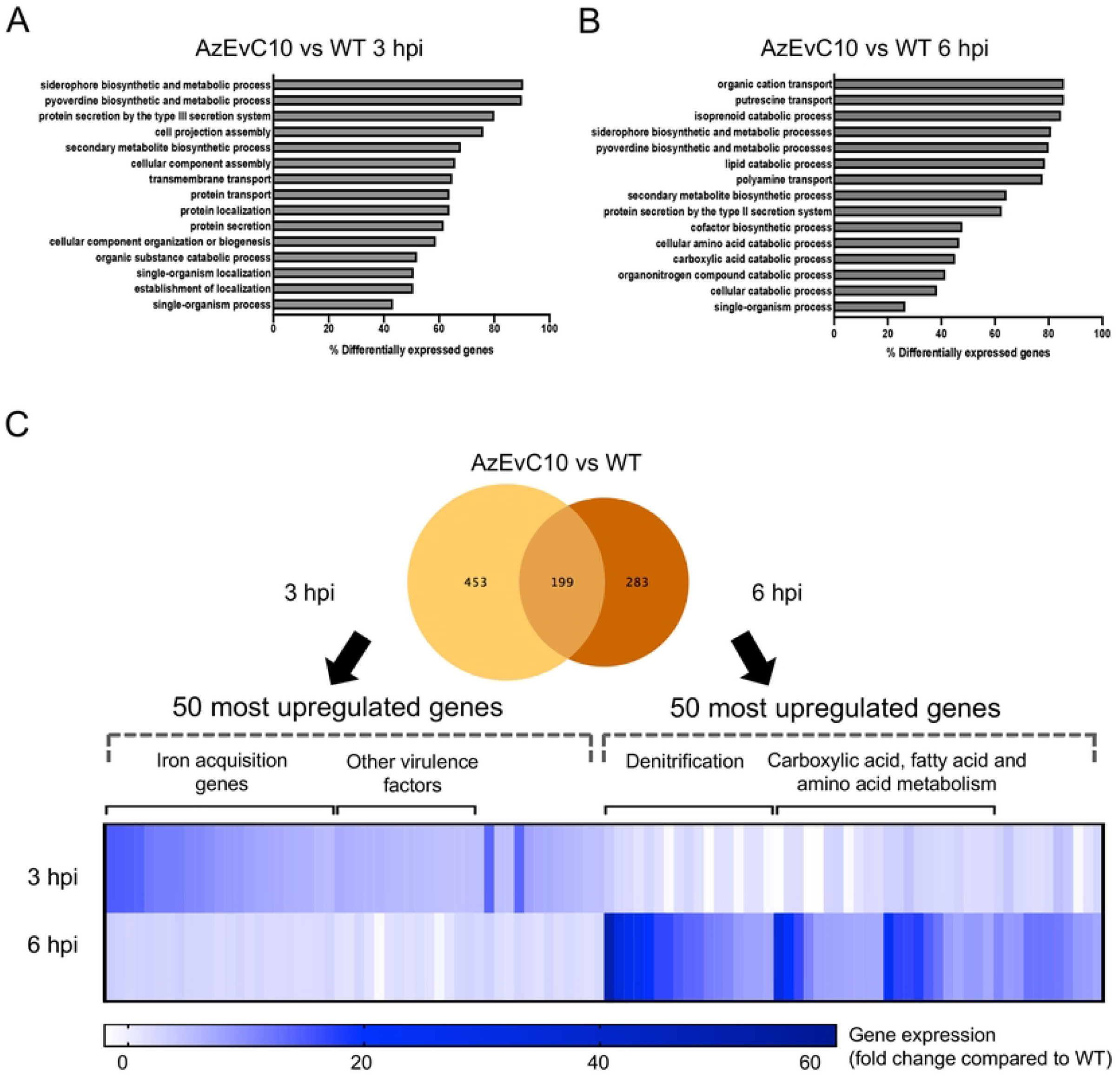
The AzEvC10 mutant upregulates virulence factor expression before metabolic gene expression during macrophage infection. **A-B**. Gene Ontology Enrichment Analyses reveal distinct enriched biological processes in the AzEvC10 mutant at 3 hpi (**A**) and 6 hpi (**B**). Percentage indicates relative number of genes in each pathway that were differentially regulated. **C**. Venn Diagram of AzEvC10 mutant DEGs compared to WT PAO1 at 3 hpi and 6 hpi of BMDM. The 50 most upregulated genes at each timepoint were selected and the expression is presented in mean fold-change compared to WT PAO1 (n=2-5 independent replicates for each experiment).

To confirm these findings, we compared AzEvC10 differentially expressed gene (DEG) sets at 3 hpi and 6 hpi and selected the 50 most upregulated genes at each timepoint (Figure 4C and Table S1). At 3 hpi, 23 (46%) of the most upregulated genes were involved in pyoverdine biogenesis and siderophore transport. Virulence factor secretion genes were also present in this gene set, with 13 (26%) upregulated genes including 10 genes involved in T3SS. At 6 hpi, 17 (34%) of the most upregulated genes were involved in denitrification processes. The other 24 (48%) genes were involved in carbohydrate, fatty acid and amino acid degradation, metabolism, and transport. These different signatures at 3 hpi and 6 hpi demonstrated how the phenotype of the AzEvC10 mutant changed during the BMDM infection. In the first hours of infection, AzEvC10 secreted pyochelin and pyoverdine siderophores as well as T3SS virulence factors and regulators. At 6 hpi, the gene expression signature shifted to a highly proliferative state and adaptation to decreased oxygen availability. Interestingly, operons involved in pyochelin biosynthesis *pchABCD* and *pchEFG*, as well as the receptor *fptA* were equally overexpressed at both 3 hpi and 6 hpi (Table S1). These results support the hypothesis that the AzEvC10 mutant increased proliferation during BMDM infection due to a higher capacity to capture iron and by inducing more damage to the host cells by virulence factor secretion.

### Overexpression of siderophores results in ferroptosis in macrophages

Macrophages play an essential role in iron regulation and homeostasis. During bacterial infection, macrophages use different mechanisms to sequester iron and restrict its use by bacteria thereby limiting their pathogenicity ^29^. However, iron overload can have deleterious effects on macrophages and impair their bactericidal functions. To determine if there was iron dysregulation in macrophages infected by the AzEvC10 mutant, we quantified iron content in BMDM using Prussian blue staining (Figure 5A, 5D). The iron content was significantly greater in BMDM infected with AzEvC10 compared to those infected with the WT strain at both 3 hpi (*p*<0.01) and 6 hpi (*p*<0.05) (Figure 5A). As the iron content in BMDM infected with WT PAO1 remained the same between 3 hpi and 6 hpi, it significantly increased between 3 hpi and 6 hpi in BMDM infected with the AzEvC10 mutant (*p*<0.05).

**Figure 5.**
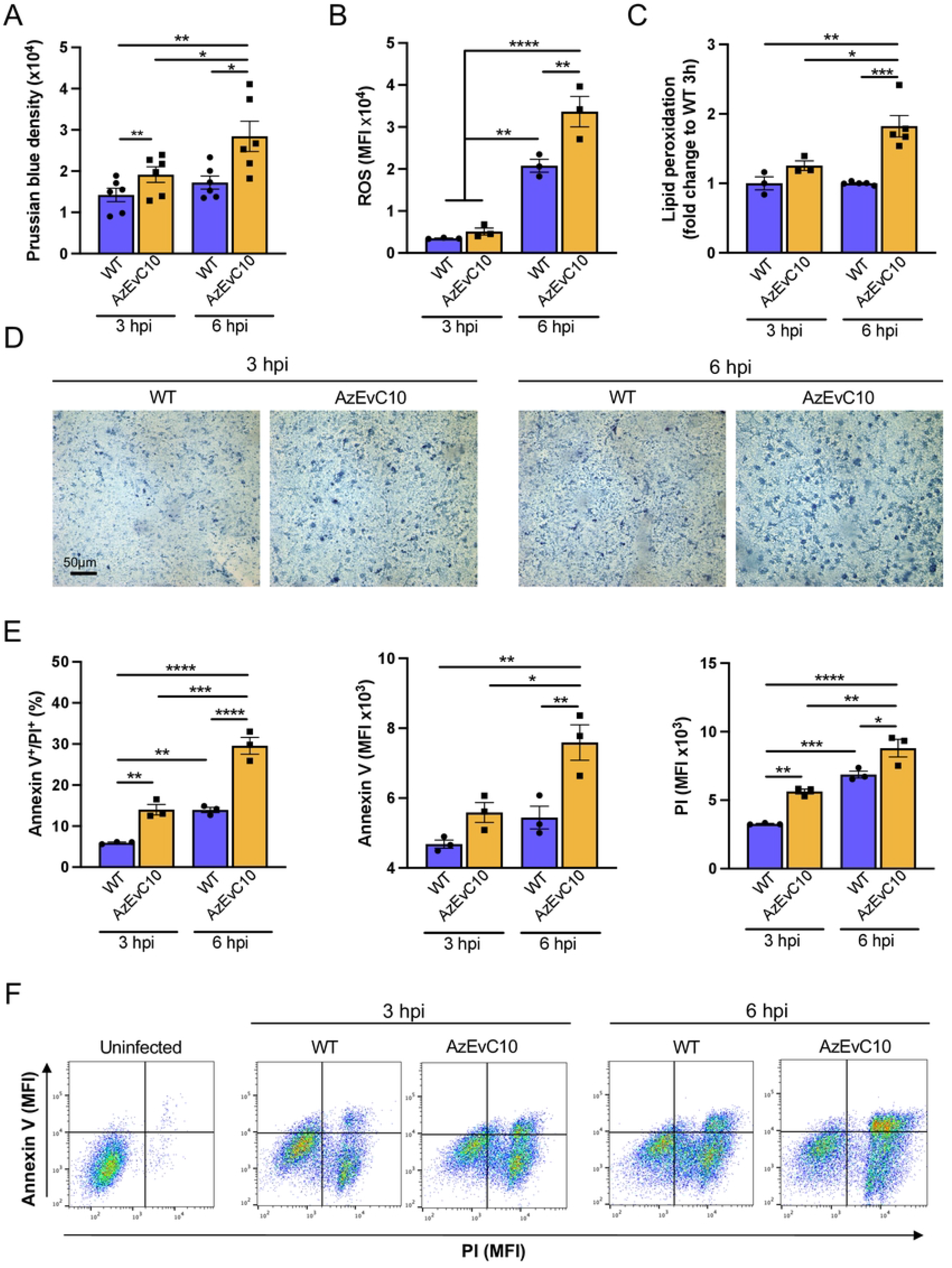
The AzEvC10 mutant induces iron dysregulation, oxidative stress, and ferroptosis in macrophages. BMDM cells were infected with WT PAO1 or AzEvC10 mutant at MOI:100 for 3 h and 6 h. **A**,**D**. Prussian blue staining of infected BMDM. Analysis was matched for experimental repeat. **B**. Quantification of ROS in infected BMDM by flow cytometry. Quantification is presented in mean fluorescence intensity (MFI). Analysis was matched for experimental repeat. **C**. Quantification of lipid peroxidation in BMDM by flow cytometry presented by the ratio of red (Ex561/Em582)/green (Ex488/Em525) fluorescence intensities and normalized to WT PAO1 3 h. **E-F**. BMDM cell death was measure by flow cytometry and characterized by BMDM cells double positive for annexin V and PI, Annexin V MFI, and PI MFI. n=3-6 independent replicates for each experiment. \**p*<0.05, \*\**p*<0.01, \*\*\**p*<0.001, #x002A;\*\*\**p*<0.0001. See Table S5 for statistical tests used and exact *p*-values.

Iron overload predisposes cells to oxidative stress through the Fenton reaction ^30–32^. Reactive oxygen species (ROS) production is a defensive mechanism used by host cells to kill intracellular pathogens. However, too much ROS can cause host damage and lead to cell death. Therefore, we quantified general ROS production in BMDM during bacterial infection (Figure 5B). As expected, both WT and AzEvC10 infected BMDM increased their production of ROS at 6 hpi (*p*<0.01 and *p*<0.0001 compared to 3 hpi respectively). However, BMDM infected with the AzEvC10 mutant had significantly greater ROS production at 6 hpi compared to macrophages infected with WT PAO1 (*p*<0.01).

We then assessed if this degree of ROS stimulated by infection with the AzEvC10 mutant was sufficient to induce cell damage. One of the consequences of iron-induced ROS toxicity in cells is lipid peroxidation, in which free radicals attack double bonds of fatty acids leading to oxidative damage of polyunsaturated fatty acids and cell structure damage ^30^. We used a ratiometric sensor to measure lipid peroxidation in BMDM and found that lipid peroxidation was significantly greater in AzEvC10 infected BMDM compared to those infected with WT PAO1 (p<0.05) (Figure 5C). Iron overload, ROS production, and lipid peroxidation can lead to ferroptosis, a form of iron-mediated programmed cell death. Consistent with this, we found that BMDM infected with AzEvC10 exhibited high levels of annexin V and propidium iodide staining (Figure 5E). The increase of these programmed cell death markers matched the increases in AzEvC10-mediated macrophage death shown in Figure 1B. It is possible that other programmed or non-programmed cell death pathways (pyroptosis, necroptosis, necrosis, etc.) are also involved in BMDM death, although together our results suggest that the AzEvC10 mutant impairs iron regulation in macrophages and induces excessive ROS and lipid peroxidation leading to ferroptosis of macrophages.

### Gallium treatment efficiently kills multidrug-resistant and hypervirulent *P. aeruginosa*

Gallium is an ion with similar properties to iron and has been proposed as an antimicrobial agent against *P. aeruginosa* ^33^. Because virulence of the AzEvC10 mutant included a higher capacity to acquire iron and induce toxicity with siderophores, we sought to test if gallium was efficient against this multidrug-resistant mutant during BMDM infection. We first confirmed inhibition of bacterial growth with 150 μM gallium in LB broth (Figure 6A). We then tested gallium treatment during BMDM infection (Figure 6B-6F). As shown above (Figure 1B), the AzEvC10 mutant was markedly more toxic to macrophages than WT PAO1 (Figure 6B). Although gallium significantly decreased BMDM cytotoxicity caused by either strain in a dose-dependent manner (Figure 6B), we saw a significant difference in the protective effectiveness of this iron-like ion. Whereas cytotoxicity caused by WT PAO1 was completely abolished with the lowest dose of gallium (100 µM), BMDM death caused by AzEvC10 remained significantly elevated at all doses. Interestingly, at 750 µM gallium pyochelin levels were significantly decreased and pyoverdine production was abolished in the AzEvC10 mutant without impacting bacterial growth at 3 hpi (Figure 6C). Siderophore production was still inhibited at 6 hpi but was accompanied by significant inhibition (∼2.5-3.5-fold) of bacterial growth (Figure 6D).

**Figure 6.**
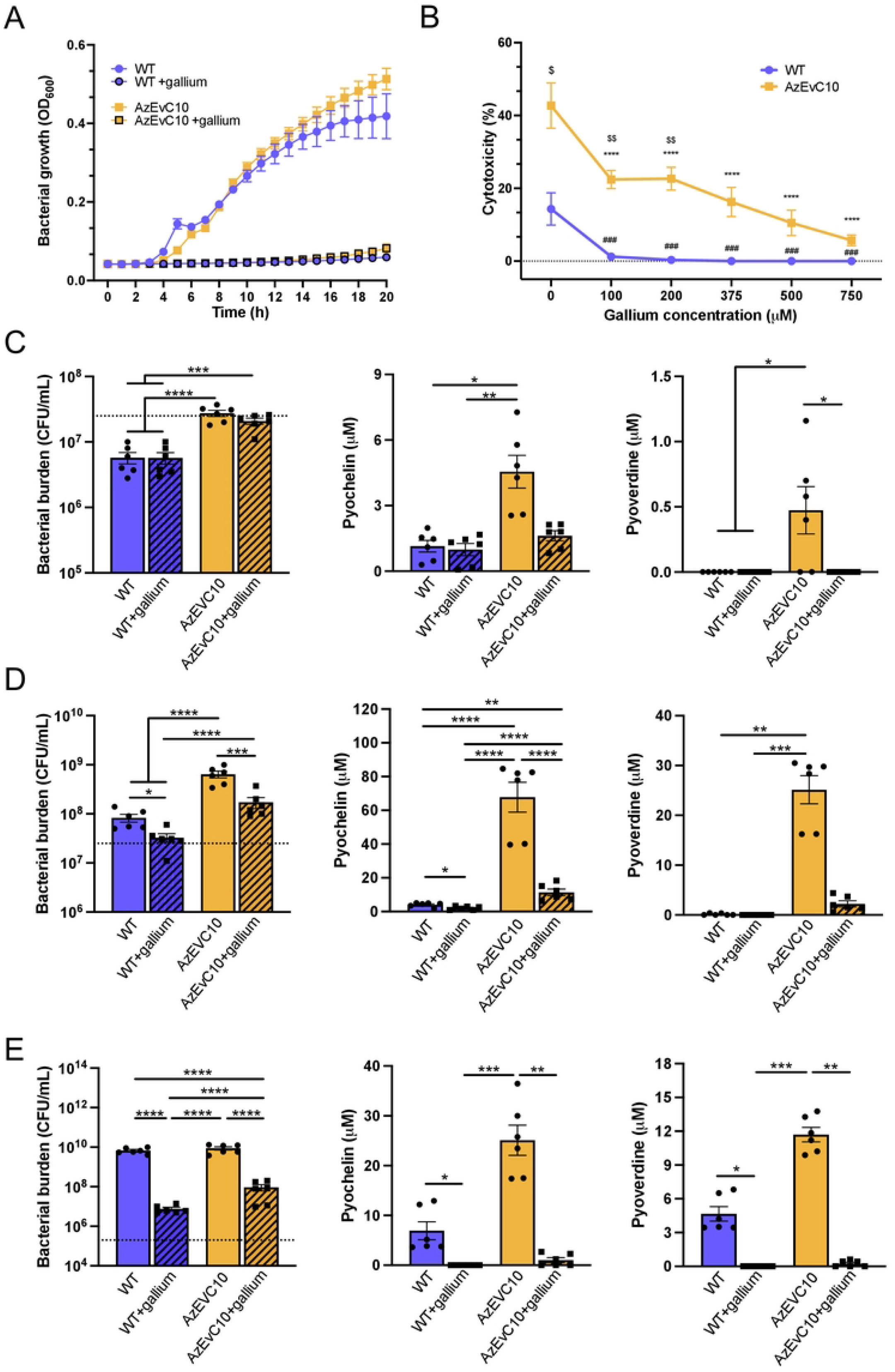
Gallium efficiently inhibits AzEvC10 siderophore production and proliferation during macrophage infection. **A**. WT PAO1 and AzEvC10 mutant growth curves in LB broth in the presence or absence of 150 μM gallium. **B**. BMDM were infected with either WT PAO1 or the AzEvC10 mutant (MOI:100) and treated with different concentrations of gallium for 6 h. BMDM cytotoxicity assessed by LDH assay: #represents the comparisons between WT with gallium to the WT no gallium baseline; *represents the comparisons between AzEvC10 with gallium to AzEvC10 no gallium baseline; $represents the comparisons between AzEvC10 and WT at the same gallium concentration. **C-D**. BMDM were infected with either WT PAO1 or the AzEvC10 mutant (MOI:100) and treated with 750 μM gallium for 3 h **(C)** and 6 h **(D)**. Bacterial burden was determined by viable CFU plate counts. Dotted lines represent the initial infection inoculum, 2.5 × 10^7^ CFU/mL. **E**. BMDM were infected with either WT PAO1 or AzEvC10 mutant (MOI:1) and treated with 750μM gallium for 24 h. Bacterial burden was determined by viable CFU plate counts. Dotted lines represent the initial infection inoculum, 2.5 × 10^5^ CFU/mL. n=6 independent replicates for each experiment. \**p*<0.05, \*\**p*<0.01, \*\*\**p*<0.001, \*\*\*\**p*<0.0001. See Table S5 for statistical tests used and exact *p*-values.

We then infected BMDM for 24 h using a lower MOI (MOI:1) with or without gallium. Gallium was efficient at killing both WT (∼1000-fold decrease, *p*<0.001) and AzEvC10 mutant (∼100-fold decrease, *p*<0.0001) *P. aeruginosa* (Figure 6E). As shown at the higher MOI, production of pyochelin and pyoverdine was greater in the AzEvC10 mutant compared to WT PAO1 (*p*<0.0001) and both were abolished with gallium at 24 hpi (Figure 6E). Our results suggest that gallium has bactericidal potential against multidrug-resistant and hypervirulent *P. aeruginosa* strains, including strains like the AzEvC10 mutant.

### Virulence and metabolic adaptation are common features of pulmonary infections

*P. aeruginosa* is a versatile bacterium that adapts to different environments. While the *in vitro* BMDM infections recapitulate interactions between *P. aeruginosa* and macrophages during *in vivo* infections, this model lacks several key environmental characteristics that mirror real-world human infections. These missing features include other host cells like neutrophils and epithelial cells, as well as the physical and chemical properties of sputum in CF airways. Thus, gene expression measured during *in vitro* experiments may not always accurately predict bacterial functioning during clinical infections.

To test if the pathoadaptive phenotype observed in the AzEvC10 mutant during macrophage infection was reflective of *P. aeruginosa* gene expression profiles during real-world infections, we analyzed RNA-sequencing data from CF sputum and subsequently isolated *P. aeruginosa* clinical isolates grown *in vitro* ^34^ (Figure 7A). In accordance with published findings ^34^, our analyses showed that pyochelin biosynthesis enzymes (*pchABCD, pchEFG*) and receptors (*fptA, fptB*) were significantly overexpressed during pulmonary infections of CF patients but not when grown *in vitro* (Figure 7B, Table S2). We then compared the genes that were differentially expressed between the sputum samples and *in vitro* grown *P. aeruginosa* with the DEGs of our WT PAO1 or AzEvC10 strains between macrophage infection and LB growth. As expected, common upregulated genes between AzEvC10 mutant 3 hpi and sputum included pyoverdine and pyochelin biosynthesis enzymes and transport receptors (Figure 7C, Table S2). Other common genes were involved in the response to oxidative stress (*ahpC*), T1SS (*aprA, aprFED*), and regulation of the T3SS (*rsmZ* and *pcrG*, respectively). Interestingly, the alkaline protease (*aprA*) and its secretory apparatus (*aprFED*) expressions are regulated by iron levels and facilitate iron acquisition by the proteolytic cleavage of transferrins^35,36^. This highlights the importance of iron acquisition during infection and a role for *rne* mutants in iron-mediated bacterial virulence. A subset of commonly upregulated genes shared by the AzEvC10 mutant 6 hpi and sputum *P. aeruginosa* are known to be expressed under anaerobic conditions, including the *nir* and *nor* operons, cytochrome C oxidase subunits (*ccoP2* and *ccoQ2*), arginine deaminase (*arcA*), and malate/L-lactate dehydrogenase (*dpkA*) (Figure 7D, Table S2). This switch to anaerobic metabolism contributes to increased fitness of *P. aeruginosa* under hypoxic conditions. The open reading frame PA3271 was also overexpressed in both AzEvC10 at 6 hpi and sputum and encodes a probable two-component sensor. Because more than 50% of two-component systems have been implicated in virulence and *in vivo* fitness and colonization ability ^15^, this suggests that the sensor may be important for *P. aeruginosa* responses to macrophages. Hence, our results show that the pathoadaptive changes seen in AzEvC10 during macrophage infection share similarities with *P. aeruginosa* behavior during pulmonary infections.

**Figure 7.**
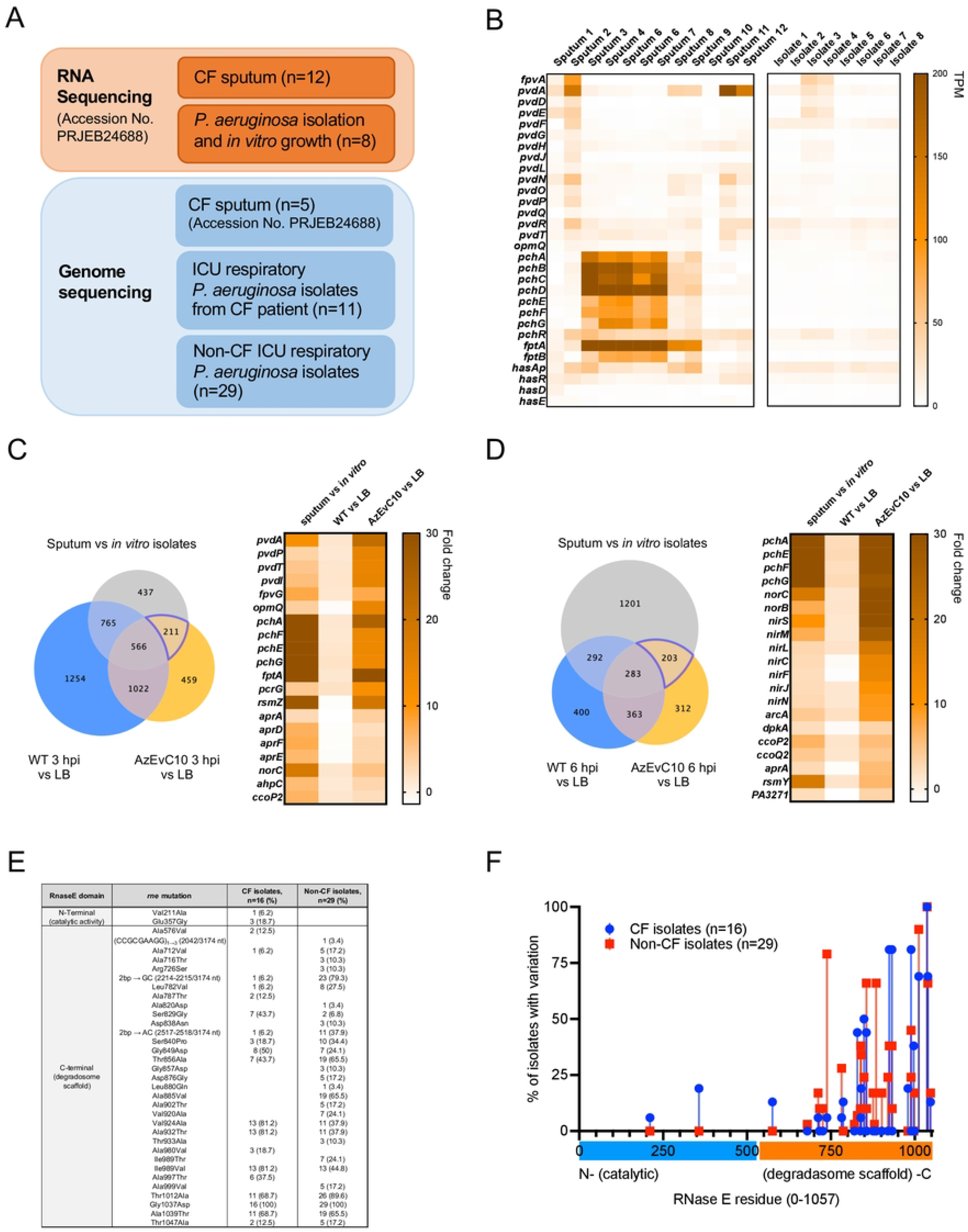
Virulence and metabolic adaptation are common features of *P. aeruginosa* pulmonary infections. **A**. Schematic representation of the experimental design for RNA and whole genome sequencing of *P. aeruginosa* clinical isolates. **B**. Individual expression of iron acquisition genes in CF sputum (n=12) and *in vitro* grown clinical isolates (n=8) expressed in normalized number of reads (TPM) **C-D**. Expression of selected iron and metabolic genes commonly unregulated by clinical isolates in sputum and by the AzEvC10 mutant during BMDM infection at 3 hpi **(C)** and 6 hpi **(D)** (n=2-5). **E**. List of non-synonymous mutations detected in the *rne* gene in *P. aeruginosa* clinical isolates. **F**. Representation of the genomic location of *rne* mutations in clinical isolates. Fisher’s exact test was used to compare the prevalence of mutations in amino-terminal vs carboxy-terminal halves (*p*<0.0001) (Table S5).

### Clinical isolates evolve multiple mutations in *rne* gene

The *rne* gene encodes RNase E endonuclease, a 1,057-residue protein involved in RNA processing and decay ^37^. Because the virulent phenotype of AzEvC10 mutant is caused by a 50-bp deletion in the of 3’ end of the *rne* gene encoding the enzyme’s carboxy-terminus, we assessed if mutations in this region were present in human clinical isolates. We performed whole genome sequencing on *P. aeruginosa* isolates from both CF (n=16) and non-CF (n=29) respiratory infections (Figure 7A). We found that all isolates (n=45) had multiple non-synonymous mutations in the *rne* gene (Figure 7E). All non-synonymous mutations except for two were found in the second half of the gene, which correspond to the carboxy-terminal half of the RNase E protein (*p*<0.0001) (Figure 7F). Based on our observations in the AzEvC10 mutant during macrophage infection, these results suggest that *rne* variations evolved in *P. aeruginosa* clinical isolates may have contributed to their colonization and/or fitness during pulmonary infections. However, because clinical isolates likely have additional mutations in other genes, it is difficult to directly demonstrate the role of *rne* mutations on the gene expression and fitness of these clinical isolates. Further investigation will be needed to determine which variants affect RNase E functions and bacterial phenotypes during infection.

## Discussion

*P. aeruginosa* is present in more than 50% of adults with CF and is thought to adapt to the CF lung environment by decreasing production of virulence factors and acquiring antibiotic resistance features. In this study, we challenge this concept of an inverse correlation between acquisition of antibiotic resistance and virulence factor production. In a previous study, we showed that aztreonam selected for mutations of the MexAB-OprM regulator genes, including *nalD* ^26^. Mutants carrying mutations in the *nalD* gene were multidrug resistant and found in 7-33% of clinical isolates, depending on the study ^26,38–42^. In this study, we showed that the multidrug-resistant AzEvC10 mutant evolved a compensatory mutation in the *rne* gene, which led to increased survival and increased proliferation during macrophage infection compared to WT and induced significantly higher cytotoxicity in macrophages (Figure 1). Increased macrophage killing *in vivo* could lead to impaired host protection against pathogens and exacerbated inflammation ^4,5,43–47^. We showed that although aztreonam resistance was due to a loss-of-function *nalD* mutation, the AzEvC10 hypervirulent phenotype resulted from a *rne* gene variant (Figure 2, Figure S2). The evolution of increased virulence towards macrophages was surprising, given that this phenotype evolved only in response to aztreonam, suggesting that antibiotic selection alone can lead to hypervirulent phenotypes.

RNase E is an endonuclease with two main domains: a catalytic amino-terminus, and a carboxy-terminus which serves as a scaffold for the degradosome complex ^48,49^. In *E. coli* the last residues of C-terminus contain the binding site for the exoribonuclease Polynucleotide phosphorylase (PNPase) ^49^. Thus, a 17-residue deletion in that region as seen in our AzEvC10 could prevent or impair PNPase binding and overall degradosome activity. Although the absence of the degradosome complex enhanced RNA half-lives in *E. coli*, some studies suggested that PNPase was unnecessary for RNA degradation and was rather a scavenger of RNA intermediates ^37,50–52^. However, mutants lacking PNPase or the PNPase-binding site in RNase E were sufficient to increase RNA half-lives of specific mRNAs, suggesting that the presence of PNPase in the degradosome complex is necessary for degradation of specific mRNAs ^53^. In our study, a 17-residue deletion in the PNPase-binding site of RNase E was sufficient to create a global transcriptomic shift in the AzEvC10 mutant during macrophage infection. We found a striking upregulation of pyoverdine and pyochelin secretion by AzEvC10, *rne*_Δ50bp_, and *rne*::Tn strains (Figure 3, Table S1). Interestingly, heme sensor genes of the Has system were also upregulated, but not those involved in the Phu system (Figure 3, Table S1). This suggests that the mutation in AzEvC10 RNase E impacts specific substrates as opposed to the whole mRNA pool. *E. coli* RNase E was also shown to interact with RNA-binding proteins such as Hfq in a degradosome-independent manner ^51,54,55^. However, the literature differs on the necessity of this interaction for RNA cleavage ^51,54,56^. The RNase E carboxy-terminal region was also shown to be essential for binding and localization of the degradosome to the bacterial membrane, which could affect its activity ^57,58^. In *P. aeruginosa* in general, and in our study, it is unknown whether the RNase E-Hfq interaction or cellular localization are impaired in the AzEvC10 mutant. However, due to the location of the AzEvC10 RNase E mutation, we believe this to be unlikely. Ultimately, future work will be necessary to determine the precise effects of the RNase E C-terminal variation on its biochemical functions.

The upregulation of iron acquisition pathways preceded the metabolic changes (Figure 4, Table S1), suggesting that this increased virulence and capacity to capture iron drove the proliferative phenotype of AzEvC10 mutant during macrophage infection. Siderophores can also be secreted for other purposes than iron acquisition. For instance, pyochelin can increase *P. aeruginosa* competitiveness against other bacteria and cause tissue and host cell injury by catalyzing the formation of ROS via the Fenton reaction. ^59–61^. On the other side, bacteria protect themselves from ROS by upregulating enzymes involved in superoxide metabolism (*ahpC, sodM*) (Table S1). The T3SS can also induce macrophage lysis ^9,10^ and was previously shown to be positively regulated by RNase E in *P. aeruginosa* ^62^. Although T3SS genes were upregulated at 3 hpi by the AzEvC10 mutant, their negative regulators, the *rsmY* and *rsmZ* small non-coding RNAs were upregulated as well (Figure 4, Table S1). Therefore, it is not clear whether the T3SS is active or inactive in the AzEvC10 strain. Future studies will examine the role of this virulence system in the host cell damage and bacterial escape using different *rne* mutants. RNase E or homologs were also involved in bacterial virulence and metabolic adaption under stress conditions in different bacterial species ^51,55,63,64^. We speculate that the stress induced by phagocytosis triggers the secretion of virulence factors by the bacteria to survive the phagocytic process and eliminate a threat.

The AzEvC10 mutant significantly increased cell toxicity by disrupting iron regulation and inducing excessive ROS in macrophages, lipid peroxidation, and ferroptosis (Figure 5). While other cell death pathways, like pyroptosis or necroptosis, may also be involved in the BMDM cell death, the lipid peroxidation is a hallmark of ferroptosis ^65^. This was somewhat unexpected, as macrophages activated with LPS express iNOS to resist ferroptosis triggered by chemical treatment with the pro-ferroptotic GPX4 inhibitor, RSL3 (1S,3R)-2-(2-chloroacetyl)-2,3,4,9-tetra-hydro-1-[4-(methoxycarbonyl)phenyl]-1H-pyrido[3,4-b]indole-3-carboxylic acid)^66^. However, in RNA-seq analyses we observed no decrease in iNOS expression (not shown) in the mutant infected BMDM, yet lipid peroxidation was still prevalent, indicating that activated macrophages may be susceptible to ferroptosis during infection. This apparent discrepancy between the mutant infected BMDM and LPS activated BMDM could be explained by the competition between the host and microbe for iron. Macrophages respond to bacterial infection by sequestering iron, which prevents its use by pathogens. They can also steal iron directly from pyochelin and pyoverdine siderophores ^67^. Increased iron uptake by BMDM from the media and siderophores could explain the iron overload observed in AzEvC10-infected BMDM (Figure 5A, 5D). About 0.2-5% of cellular iron is considered transiently mobile, promotes the formation of highly reactive ROS via the Fenton reaction, and may lead to lipid peroxidation ^30,31^, which together could explain the increased ferroptosis of the AzEvC10-infected BMDM. Furthermore, it was shown that ferripyochelin was an efficient catalyst for hydroxyl radical formation and induced cell injury ^60,61^. Pyoverdine was also recently shown to translocate into host cells and disrupt iron mitochondrial homeostasis ^68^. Although these studies differ on which siderophore causes more damage to cell host, they undoubtedly support a role for siderophores in host cell toxicity.

Lipoxygenases and cyclooxygenases can also oxidize lipids. It was recently demonstrated that some biofilm-producing *P. aeruginosa* isolates overexpressed the enzyme pLoxA, which induced ferroptosis in human bronchial epithelial cells by oxidation of arachidonic acid– phosphatidylethanolamines ^69^. In our study, pLoxA was not increased in the AzEvC10 mutant. Although this enzyme is involved in cell lipid oxidation, it is unlikely the root cause of increased lipid peroxidation and subsequent ferroptosis observed in the macrophages here. Lipid peroxidation is a hallmark of CF ^70,71^ and ferroptosis was recently described in CF epithelial cells ^72–74^. Our study expands the current understanding of how *P. aeruginosa* can induce ferroptosis in CF, showing that 1) multidrug-resistant *P. aeruginosa* can induce ferroptosis in macrophages, one of the most important cell types in bacterial clearance ^4–7^, and 2) RNase E variants can confer bacterial hypervirulence by triggering host cell ferroptosis. Since both ferroptosis and dysfunctional macrophages lead to exacerbated inflammation and tissue damage, it is imperative to develop therapeutics that will target this process and protect host cells. One of such treatment could be the use of ferroptosis inhibitors to prevent cell death. Another approach, the one chosen in the current study, is to target the root cause of ferroptosis, bacterial siderophores, by using iron competitors.

Since siderophores were upregulated in the AzEvC10 mutant and induced toxicity during macrophage infection, we sought to target and inhibit these molecules using gallium nitrate.

Previous work suggested gallium as a novel therapy for its antimicrobial properties against *P. aeruginosa* and *Klebsiella pneumoniae* ^33,75–80^. In the present study, gallium completely abolished siderophore-mediated cytotoxicity in BMDM (Figure 6). Moreover, gallium treatment for 24 h decreased bacterial growth by a factor of 100 to 1000-fold. This is important since AzEvC10 mutant *P. aeruginosa* is resistant to multiple antibiotics currently used in CF patients ^26^. A first clinical trial conducted by Goss *et al*. showed gallium to be a suitable treatment for CF patients ^33^. However, a larger study (NCT02354859) showed less promising results, although gallium did slightly improve respiratory symptoms and decrease bacterial density in sputum cultures. Different hypotheses may explain this lack of efficiency of gallium treatment seen in CF patients. One of them is the low concentration of gallium detected in the sputum, likely caused by the intravenous administration of the treatment ^33^. To tackle this issue, inhalation of gallium was suggested as a potential more effective mode of administration compared to intravenous delivery in a pre-clinical model ^81^. As a result, a clinical study is currently ongoing to test the safety and pharmacokinetic properties of inhaled gallium in CF patients (NCT03669614). A second hypothesis is that the people treated with gallium were infected with *P. aeruginosa* that would not be affected by the gallium treatment. Our data suggest that gallium could be an effective precision treatment for people infected with *P. aeruginosa* strains that overexpress siderophores. Finally, the experimental design used in our study is a model of acute infection. This leads us to believe that gallium treatments could be more effective in acute rather than chronic respiratory infections by *P. aeruginosa*.Increased iron acquisition, resistance to oxidative stress and changes in metabolism were detected in *P. aeruginosa* in sputum and in the AzEvC10 mutant during macrophage infection (Figure 7). These differentially expressed pathways were previously described in a very elegant study by Rossi *et al*. ^34^. The comparison between our experimental model and clinical isolates from CF sputum support our findings by showing that phenotypes detected in the AzEvC10 mutant are also detected during infection. Furthermore, we found a striking abundance of variations in the carboxy-terminus of the *rne* gene in *P. aeruginosa* clinical isolates (Figure 7), suggesting that *rne* variations evolved in *P. aeruginosa* clinical isolates could enhance their colonization and/or fitness during pulmonary infections. Comparing an evolved lab strain with clinical isolates does have limitations. An obvious limitation is that the accumulation of genetic mutations in these clinical isolates during CF chronic infection could have secondary effects on the phenotypes that might be induced by RNase E variations. Clinical isolates from CF sputum were shown to have distinct genetic backgrounds ^34^. These differences could impact genes nvolved in iron acquisition, resistance to oxidative stress, or metabolic pathways. Therefore, the direct effects of *rne* mutations on these pathways is difficult to directly demonstrate without in-depth future investigation.

Some limitations exist in our study. We speculated that the degradosome assembly and/or function was impaired in the AzEvC10 mutant, explaining its increased virulence by siderophore overexpression. However, have not directly tested the effects of the *rne* mutation on degradosome function, nor the interaction of RNase E with PNPase. The mutation could also impair RNase E interaction with other proteins, like RNA-binding proteins, rather than PNPase. These aspects will need to be elucidated in further studies. Finally, we used healthy instead of CF macrophages for infections. It was previously shown that a dysfunctional cystic fibrosis transmembrane conductance regulator (CFTR) channel sensitized epithelial cells to lipid peroxidation and ferroptosis ^72^. However, CFTR corrector only partially reversed lipid peroxidation ^74^. Furthermore, *P. aeruginosa* equally caused lipid peroxidation and ferroptosis in the airways of CFTR^+/+^ and CFTR^-/-^ mice and in human airway epithelial cells expressing WT CFTR or F508Del mutant CFTR ^73^. Thus, it is unclear whether *P. aeruginosa* overexpressing siderophores would display a greater toxicity to macrophages expressing a dysfunctional CFTR channel.

Our study underscores how a second-site compensatory mutation evolved during antibiotic selection can restore virulence in multidrug-resistant *P. aeruginosa*. Furthermore, we demonstrate a role for RNase E endonuclease in bacterial virulence and metabolic adaptation during infection. We also showed that mutations in *rne* genes are prevalent in clinical isolates from acute and chronic pulmonary infections. Second-site mutations, including mutations in *rne* genes, could be a key mechanism that lead multidrug- and extensively drug-resistant *P. aeruginosa* to be hypervirulent in CF. Because infections with resistant *P. aeruginosa* are increasing worldwide ^20–22^, the need for new therapies is more urgent than ever. Gallium nitrate is a promising therapy for these “superbugs” and targeting ferroptosis may represent another means to reduce *P. aeruginosa* pathogenesis.

## Author Contributions

Conceptualization, M.V. and P.J.; methodology, M.V. and P.J.; formal analysis, M.V., A.C.M.G., S.P.L., D.C., E.D., M.B., C.B., J.S.L., W.C.P. and P.J.; investigation, M.V., A.C.M.G., S.P.L., D.C., E.D., M.B., C.B., J.S.L., W.C.P. and P.J.; resources, Y.D., J.S.L., W.C.P., and P.J.; data curation, M.V., A.C.M.G., S.P.L., D.C., E.D., M.B., C.B., J.S.L., W.C.P. and P.J.; writing—original draft preparation, M.V. and P.J.; writing—review and editing, M.V., A.C.M.G., S.P.L., D.C., E.D., M.B., C.B., J.S.L., W.C.P. and P.J.; supervision, M.V. and P.J.; project administration, M.V. and P.J.; funding acquisition, W.C.P., J.S.L., and P.J. All authors have read and agreed to the published version of the manuscript.

## Funding

This research was funded by grant numbers JORTH17F5 and JORTH19P0 from the Cystic Fibrosis Foundation, grant numbers K22AI127473 and R01AI14642 from the NIH/National Institute of Allergy and Infectious Diseases, R01Hl136143 from the NIH/National Heart, Lung, and Blood Institute, as well as a sub-award from grant number UL1TR001881 from the NIH/National Center for Advancing Translational Science (NCATS) UCLA CTSI.

## Acknowledgments

We would like to thank members of the Jorth Lab and Holly Huse for helpful discussions and feedback on this manuscript. We are grateful to Pradeep K. Singh, Colin Manoil, and Peter Chen for the generous gifts of cells and strains used in this study. We also thank Warren G. Tourtellotte for the use of his Zeiss Apotome fluorescence microscope. We are grateful to the Applied Genomics, Computation and Translational Core at Cedars Sinai Medical Center for their help with the RNA sequencing. Finally, we warmly thank the whole Pulmonary Translational Research Core team at University of Pittsburgh for the *P. aeruginosa* respiratory infection isolates used in this study.

## Declaration of Interests

The authors declare no conflict of interest.

## Supplemental Figure Legend

**Figure S1. Susceptibility to BMDM growth medium and amikacin**

**A**. Bacterial survival determined by viable CFU plate counts after inoculating BMDM growth medium with *P. aeruginosa* strains in the absence of BMDM. Dotted line represents the initial inoculum, 2.5 × 10^7^ CFU/mL. **B**. BMDM growth media was inoculated with WT PAO1 and AzEvC10 mutant with or without amikacin 50, 100 or 200 μg/mL and incubated for 60 min at 37°C and 5% CO_2_. Dotted line represents the initial inoculum, 2.5 × 10^7^ CFU/mL. No CFU was recovered at 200 μg/mL for both WT PAO1 and AzEvC10 mutant, and this concentration was used for assays to kill extracellular bacteria during BMDM infection. n=3-6 independent replicates for each experiment. \*\**p*<0.01. See Table S5 for statistical tests used and exact *p*-values.

**Figure S2. Virulence of the AzEvC10 mutant is not reversed by *nalD* complementation or *mexAB* efflux pump deletion**

**A**. Genome diagram showing coverage of sequencing reads aligning to the *nalD* gene in WT PAO1 and the AzEvC10 mutant. The single base substitution leading to *nalD*_T158P_ in AzEvC10 mutant is highlighted in red. **B-C**. *nalD* complementation in AzEvC10 mutant was confirmed by RT-qPCR **(B)** and aztreonam MIC **(C). D-E**. BMDM were infected with either WT PAO1, AzEvC10, or AzEvC10 carrying pMQ72::*nalD*_WT_ at MOI:100 for 6h. **D**. Bacterial burden determined by viable CFU plate counts. Dotted line represents the initial infection inoculum, 2.5 × 10^7^ CFU/mL. **E**. BMDM cytotoxicity was assessed by LDH assay. **F**. Gel electrophoresis showing *mexAB* deletion in WT PAO1 and AzEvC10 strains. **G**. Aztreonam MIC was by Etest strip. **H-I**. BMDM were infected with either WT PAO1, AzEvC10 or the strains containing the *mexAB* deletion at MOI:100 for 6h. **H**. Bacterial burden determined by viable CFU plate counts. Dotted line represents the initial infection inoculum, 2.5 × 10^7^ CFU/mL. **I**. BMDM cytotoxicity was assessed by LDH assay. n=3-4 independent replicates for each experiment. \**p*<0.05, \*\**p*<0.01, and \*\*\**p*<0.001. See Table S5 for statistical tests used and exact *p*-values.

## Supplemental Tables

**Table S1. 50 most upregulated genes in AzEvC10 during macrophage infection**

**Table S2. Common DEGs between sputum and AzEvC10 during macrophage infection**

**Table S3. List of strains**

**Table S4. List of primers**

**Table S5. Statistics**

## Materials and Methods

### Bacterial strains and growth conditions

Bacterial strains and plasmids are listed in Table S3. *P. aeruginosa* PAO1 AzEvC10 and AzEvC10 pMQ72::*nalD* mutants were obtained from Pradeep K. Singh ^26^. The *rne* Tn mutant (*rne*::Tn) was obtained from Colin Manoil’s laboratory at the University of Washington and corresponds to the PW5993 mutant containing the genotype PA2976H05:: IS*phoA*/hah ^28^. This mutant was grown from freezer stock on Luria-Bertani (LB) agar with 10mg/mL tetracycline. All strains were maintained at 37°C in Luria-Bertani (LB) broth unless otherwise specified. Where necessary, gentamicin (Gm) was added to growth media at the following concentrations: 10 μg/mL for *E. coli* and 30 μg/mL for *P. aeruginosa*.

### *P. aeruginosa* clinical isolates

*P. aeruginosa* clinical isolates from CF (n=11) or non-CF (n=29) ICU patients with pulmonary infection were a generous gift from the Pulmonary Translational Research Core at University of Pittsburgh.

### Mutant constructions

Gene deletion mutants listed in Table S3 were generated with suicide plasmids as described previously ^82^. Suicide plasmids carrying *mexAB* deletion, *nalD*_T158P_ point mutation, 50bp deletion in *rne* gene, or WT *rne* gene were constructed in a similar way. First, two PCR fragments were generated from chromosomal DNA using corresponding primer pairs to the up- and down-stream regions of each gene (see Table S4 for primer nucleotide sequences). The two fragments were then assembled into the suicide vector pEX18Gm between using NEBuilder® HiFi DNA Assembly Cloning Kit (cat# E5520). Suicide plasmids were then transformed into *E. coli* DH5α according to the NEB protocol and verified by Sanger sequencing. *P. aeruginosa* strains were transformed by electroporation as published previously ^83^ and incubated on LB + Gm 30 μg/mL at 37°C overnight. For counter selection, isolated clones were streaked on low-salt LB containing 15% sucrose as published ^82^ and incubated at room temperature for 48 h. Gene deletions were confirmed by PCR using primers listed in Table S4 and whole genome sequencing.

### DNA extraction, purification and PCR

Plasmid DNA was prepared using a Monarch® Plasmid Miniprep Kit (NEB, cat# T1010). Genomic DNA was prepared using the DNeasy Blood & Tissue Kit (Qiagen, cat# 69504). When necessary, cDNA was purified using the Monarch® PCR & DNA Cleanup Kit (NEB, cat# T1030). PCR was performed using either KAPA HIFI 2X ready mix (KAPA Biosystem, cat# KK2602) or Expand Long Template PCR System (Sigma, cat# 11681842001). Primers used for each PCR reaction are listed in Table S4.

### Aztreonam minimal inhibitory concentration test

Etest strips were purchased from Biomerieux Vitex Inc (cat# 501758) and antimicrobial susceptibility testing was performed with the following modifications to the manufacturer’s instructions. A sterile swab was soaked in an overnight culture for each strain after growth for 18h in LB broth and excess fluid was removed by pressing it against the inside wall of the test tube. Mueller Hinton agar plates were fully streaked 4 times with the swabs. After allowing the plates to dry, an Etest gradient strip was placed in the middle of each plate. Plates were incubated at 37°C for 16–20 h and MICs were read by zone of clearing around the strip.

### BMDM isolation

Femurs and tibias from C57BL/6 mice were harvested and cleaned up with 70% ethanol. Working in a biological safety cabinet, bones were washed with 1X PBS. Bone marrow was flushed with 10 mL of BMDM growth medium (RPMI 1640 containing 20% L929 conditioned media, 10% fetal bovine serum [FBS, Omega Scientific cat# FB-11] and antibiotic-antimycotic). Antibiotic-antimycotic solution was purchased from ThermoFisher (cat# 15-240-062) and used at a final concentration of 1% (100 units/mL penicillin, 100μg/mL streptomycin, and 2.5 μg/mL fungizone). Red blood cells were lysed in 5 mL 1X RBC lysis buffer for 3 m at room temperature. Lysis was stopped by adding 30 mL of 1X PBS. Cells were pelleted, counted and plated on treated tissue culture dishes. After 24 h, non-adherent cells were plated into a petri dish and grown in BMDM growth media at 37°C and 5% CO_2_. Fresh media was added after 3 d. After 7 d, cells were plated for experiments.

### Bone marrow-derived neutrophil isolation

Femurs and tibias from C57BL/6 mice were harvested and cleaned up with 70% ethanol. Working in a biological safety cabinet, bones were washed with 1X PBS. Bone marrow was flushed with 10 mL of growth media (RPMI 1640 containing 10% FBS). The EasySep Mouse Neutrophil Enrichment Kit (StemCell, cat# 19762) was used to isolate neutrophils according to the manufacturer’s instructions. Neutrophils were used the same day of isolation.

### Mouse airway epithelial cells

Immortalized mouse airway epithelial cells (MLE-12) were a gift from Dr. Peter Chen (Cedars-Sinai Medical Center, Los Angeles). Cells were grown in Dulbecco’s Modified Eagle Medium containing 10% FBS (Omega Scientific cat# FB-11).

### L929 conditioned media

L929 mouse fibroblasts were grown in Iscove’s Modified Dulbecco’s Medium (ThermoFisher, cat# 12-440-061) with 10% FBS. Cells were plated at a density of 4 ×10^6^ cells/150 mm tissue culture treated dish and culture for 10 d at 37°C, 5% CO_2_. Medium was then collected, filtered through a 0.22 µm filter and frozen at -80°C in 50 mL aliquots.

### Cell infection with *P. aeruginosa*

BMDM, neutrophils, or AEC were plated with growth media without antibiotic-antimycotic for all experiments. Bacterial strains were streaked on LB agar and incubated at 37°C. The day before the infection, bacterial strains were grown in LB broth at 37°C overnight shaking at 250rpm. The day of infection, strains were diluted and grown to mid-exponential phase OD_600_ 0.4-0.6. Bacteria were then pelleted, washed with 1X PBS and resuspended in 1X PBS. BMDM were infected with a MOI: 100 for 3 h or 6 h, or with a MOI: 1 for 24 h. For intracellular survival experiments, BMDM were infected for 1 h, then amikacin was added to a final concentration of 200 μg/mL and left for an addition 5 h of incubation. For the escape experiments, antibiotics were added after 1 h incubation and left for 1 h. Cells were then washed with 1X PBS and fresh BMDM medium without antibiotic was added and cells were incubated for a total of 3 h or 6 h.

### Immunofluorescence

BMDM were infected with a MOI: 100 for 6 h. Cells were then washed with 1X PBS and fixed with 2% PFA for 10 m at room temperature. After 3 washes with 1X PBS, cells were permeabilized in 1X PBS + 5% Goat serum + 0.1% Triton X-100 for 5 m at room temperature. Anti-CD68 (BioRad, cat#MCA1957, 1: 250) and anti-*P. aeruginosa* (ThermoFisher, cat# MA183430, 1:500) primary antibodies were diluted in 1X PBS + 2% Goat serum. Cells were incubated with primary antibodies for 1h at room temperature, washed 3 times with 1X PBS, then incubated for 30-45 m with goat anti-mouse Alexa Fluor 555 (Invitrogen cat#A32727, 1:1000) or goat anti-rat Alexa Fluor 488 (Invitrogen cat#A11006, 1:1000) secondary antibodies. After 3 washes with 1X PBS, cells were mounted with Fluoromount G mounting media with DAPI (ThermoFisher, cat# 00-4959-52).Images were taken using a Zeiss Apotome fluorescence microscope and analyzed with Bitplane Imaris software version 9.2.1.

### Prussian blue staining

Just before use, 5% aqueous solution of hydrochloric acid was mixed 1:1 with 5% aqueous solution of potassium ferrocyanide. Fixed cells were covered with Prussian blue mixture and incubated at room temperature for 20 m. Cells were then washed multiple times with PBS +0.1% Tween20 and mounted on slides. Images were taken within 24 h using a brightfield microscope. Integrated density was quantified using a color deconvolution algorithm from Fiji software version 2.1.0^84^.

### Cytotoxicity assay

Cytotoxicity assays were performed using the Pierce LDH Cytotoxicity Assay Kit (ThermoFisher, cat# 88954) according to manufacturer’s instructions.

### Reactive Oxygen Species assay

Infected cells were washed with 1X PBS and then harvested with 1mL 5mM EDTA-1X PBS and centrifuged at 8000rpm for 1 m. Supernatants were removed, and cell pellets were washed with 1 mL FC buffer (3% FBS in 1X PBS). BMDM were resuspended in 10 μM 2′,7′-dichlorodihydrofluorescein diacetate dilution (Sigma cat# D6883) and incubated for 30 m at 37°C protected from light. Cells were then washed in 1 mL FC buffer and resuspended in FC buffer and immediately analyzed by flow cytometry (Becton Dickinson LSR Fortessa). Mean fluorescence intensity (MFI) was quantified using FlowJo software (version 10.6.1, FlowJo LLC).

### Lipid peroxidation assay

Infected cells were washed with 1X PBS and then harvested with 1 mL of 5 mM EDTA-1X PBS and centrifuged at 8000rpm for 1 m. Supernatant was removed, and cell pellets were washed with 1 mL FC buffer (3% FBS in 1X PBS). The Lipid Peroxidation Assay Kit (Abcam cat# ab243377) was used and cells were stained according to the manufacturer’s instructions. Cells were then washed in 1 mL Hank’s Balanced Salt Solution (HBSS) and resuspended in HBSS buffer and immediately analyzed by flow cytometry (Becton Dickinson LSR Fortessa). Mean fluorescence intensity (MFI) was quantified using FlowJo software (version 10.6.1, FlowJo LLC). Lipid peroxidation was quantified by calculating the red (Ex561/Em582)/green (Ex488/Em525) fluorescence ratio using FlowJo software (version 10.6.1, FlowJo LLC). Data were then normalized on WT PAO1 at 3 h.

### Cell death assay

Infected cells were washed with 1X PBS and then harvested with 1 mL of 5 mM EDTA-1X PBS and centrifuged at 8000rpm for 1 m. Supernatant was removed, and cell pellets were washed with 1 mL FC buffer (3% FBS in 1X PBS). To quantify cell death, TACS® Annexin V-FITC Apoptosis Detection Kit (R&D Systems cat# 483001K) was used and cells were stained according to the manufacturer’s instructions. Cells were then washed with the binding buffer and resuspended in binding buffer and immediately analyzed by flow cytometry (Becton Dickinson LSR Fortessa). The percentage of double positive cells and MFI were quantified using FlowJo software (version 10.6.1, FlowJo LLC).

### CFU counts to determine total and intracellular bacterial burdens

BMDM were permeabilized with 0.1% Triton X-100 for 5 m, then cells were collected using a cell scraper. For each condition, 10 μL was used to make a serial dilution and plated in duplicate on LB agar. Plates were incubated at 37°C overnight. For each condition, CFUs were counted and reported as CFUs/mL.

### Pyochelin and pyoverdine quantification

To quantify siderophores, pyochelin I & II (Toronto Research Chemicals cat# P840365) and pyoverdine (Sigma cat# P8124) were purchased and used as reference standard curves. For each condition, 100 μL of infected cell supernatant was transferred in duplicate into a black clear-bottom 96-well plate. The plate was read using a Varioskan Lux plate reader at Ex350/Em430nm (pyochelin) and Ex400/Em460nm (pyoverdine). Absolute quantifications were calculated using the pyochelin and pyoverdine standard curves.

### Growth curve in gallium

Overnight cultures were diluted to reach OD_600_∼0.005. Ten microliters of these cultures were inoculated in 96-well plates in LB broth containing different concentrations of gallium (III) nitrate hydrate (Sigma cat# 289892). Fifty microliters of mineral oil was added in each well to prevent evaporation. Plates were incubated at 37°C for 20 h, shaking for 5 s every hour and absorbance at 600 nm was recorded using a Varioskan Lux microplate reader (Thermo Fisher Scientific, cat# NC1141718).

### BMDM treatment with gallium

BMDM were infected as described above. For cytotoxicity assays, gallium was added immediately after infection to final concentrations of 100, 200, 375, 500, and 750μM for 6 h. For CFU counts, gallium was added immediately after infection to a final concentration of 750 μM for 3 h, 6 h, or 24 h.

### RNA-seq library preparation

BMDM were infected with *P. aeruginosa* strains at MOI: 100 for 3h or 6h. Cells were washed and preserved in RNAlater (Invitrogen cat# AM7021) until RNA extraction. Total BMDM and bacterial RNA was extracted using a RNeasy Mini Kit (Qiagen, cat# 74106) and resuspended in 30 μL of elution buffer. Of this volume, 25 μL was used for bacterial RNA libraries and 5 μL was used for BMDM RNA libraries. For bacterial RNA libraries, RNA from host nucleic acids and bacterial rRNA was depleted using MicrobEnrich™ Kit (Invitrogen, cat# AM1901) and MicrobExpress™ Bacterial mRNA Enrichment Kit (Invitrogen, cat# AM1905) respectively, according to manufacturer’s instructions. RNA-seq libraries were prepared for bacteria using the KAPA RNA HyperPrep Kit (KAPA BioSystem, cat# KK8540) and BMDM using the KAPA mRNA HyperPrep Kit (KAPA BioSystem, cat# KK8580), according to manufacturer’s instruction. Libraries were quantified using Quant-iT™ dsDNA Assay Kit, high sensitivity (Invitrogen, cat# Q33120) and the quality was verified on Agilent 4150 TapeStation System using D1000 High Sensitivity DNA Tape (Agilent, cat#5067-5584). Libraries were pooled to equimolar amounts and sequenced using a NextSeq 500 instrument to a depth of 400 million reads per sample (75 bp, single-ends) at the Cedars-Sinai Genomics Core Facility.

### RNA-seq data analyses

For clinical isolates, publicly available data was analyzed (Accession No. PRJEB24688) ^34^. RNA-seq data were analyzed using CLC Genomics workbench 20 (version 20.0.1). The *P. aeruginosa* PAO1 genome (Genbank accession #GCA_000006765.1) was as a reference for RNA sequencing read alignment. Differential Expression for RNA-Seq (version 2.2) analysis was performed to compare whole transcriptomes between the following groups: WT PAO1 vs AzEvC10 mutant at 3 hpi and 6 hpi, each strain during BMDM infection at 3 hpi and 6 hpi vs LB growth, and CF sputum vs *in vitro* clinical isolates. Reads were normalized in transcripts per million reads (TPM). Gene Set Test (version 1.1) using Gene Ontology (GO) biological processes annotations was performed to reveal enriched biological processes in the AzEvC10 mutant after 3 h and 6 h of BMDM infection. Venn Diagram for RNA-Seq tool (version 0.2) analysis was performed using a minimal absolute fold-change of 1.5 and false discovery rate (FDR) *p*-value of 0.05. The lists of DEGs are available in Table S1 and Table S2.

### Whole genome sequencing of clinical *P. aeruginosa* isolates

*P. aeruginosa* clinical isolates from CF (n=11) and non-CF (n=29) ICU patients were a generous gift from the Pulmonary Translational Research Core at University of Pittsburgh. These isolates were sent for whole genome sequencing to the Microbial Genome Sequencing Center (MiGs). Genome sequencing data will be made available upon publication in the NCBI Sequence Read Archive. These samples were analyzed alongside publicly available whole genome sequencing data from CF sputum (n=5) (Accession No. PRJEB24688) ^34^. Variant calling program Breseq version 0.35.5 was utilized to identify mutations in the isolates’ genomes relative to the PAO1 reference (Genbank accession #GCA_000006765.1)^85^. The identified mutations were filtered for indels and non-synonymous SNPs and then subsequently filtered to identify unique mutations found only in the *rne* gene.

### Statistical analyses

Normality and homogeneity of variance were assessed by Shapiro-Wilk and Brown-Forsythe tests, respectively. Data was log-transformed prior to analysis where necessary to meet assumptions necessary for parametric testing, else non-parametric rank testing was used.

Based on data distributions, analysis was performed for matched data with paired Student t-test or Wilcoxon matched pairs rank test; for two groups with a two-sample Student t-test or the nonparametric Mann-Whitney; for three or more groups with ANOVA followed by Tukey’s test or the non-parametric Kruskal-Wallis test with Dunn’s correction. Where data were collected across experimental repeats or repeated measures, data were considered as matched sets, and RM-ANOVA was used. Cytotoxicity dose curves were analyzed with two-way ANOVA. For all parametric testing, residuals were inspected to confirm fit. Fisher’s exact test was used to analyze categorical data. All testing was considered significant at the two-tailed p-value of <0.05. Analysis performed with GraphPad Prism v9. The *p*-values are listed in Table S5.

## Notes

### Competing Interest Statement

The authors have declared no competing interest.

